# Predicting Antimicrobial Activity for Untested Peptide-Based Drugs Using Collaborative Filtering and Link Prediction

**DOI:** 10.1101/2022.11.16.516845

**Authors:** Angela Medvedeva, Hamid Teimouri, Anatoly B. Kolomeisky

**Affiliations:** Department of Chemistry, Rice University, Houston, Texas, United States; Center for Theoretical Biological Physics, Rice University, Houston, Texas, United States; Department of Chemical and Biomolecular Engineering, Rice University, Houston, Texas, United States; Department of Physics and Astronomy, Rice University, Houston, Texas, United States

## Abstract

The increase of bacterial resistance to currently available antibiotics has underlined the urgent need to develop new antibiotic drugs. Antimicrobial peptides (AMPs), alone or in combination with other peptides and/or existing antibiotics, have emerged as promising candidates for this task. However, given that there are thousands of known AMPs and an even larger number can be synthesized, it is inefficient to comprehensively test all of them using standard wet lab experimental methods. These observations stimulated an application of machine-learning methods to identify promising AMPs. Currently, machine learning studies frequently combine very different bacteria without considering bacteria-specific features or interactions with AMPs. In addition, the sparsity of current AMP data sets of antimicrobial activity disqualifies the application of traditional machine-learning methods or renders the results unreliable. Here we present a new approach, featuring neighborhood-based collaborative filtering, to predict with high accuracy a given bacteria’s response to untested AMPs, AMP-AMP combinations, and AMP-antibiotic combinations based on similarities between bacterial responses. Furthermore, we also developed a complementary bacteria-specific link approach that can be used to visualize networks of AMP-antibiotic combinations, enabling us to suggest new combinations that are likely to be effective. Our theoretical analysis of AMP physico-chemical features suggests that there is an optimal similarity between two different AMPs that exhibit strong synergistic behavior. This principle, alongside with our specific results, can be applied to find or design effective AMP-AMP combinations that are target-specific.

**Author summary:** It is well known that combinations of different antimicrobial peptides (AMPs), in comparison to single AMP species, can lead to more efficient antimicrobial activity, but the large number of possible combinations requires the application of efficient machine-learning methods. We developed a new approach consisting of collaborative filtering, link prediction, and AMP feature analysis to predict previously-unknown, bacteria-specific activity of AMP combinations, suggest novel synergistic AMP-antibiotic combinations, and guide future design of effective AMP-AMP combinations.

## Introduction

Bacterial resistance to conventional antibiotics is becoming a major global health threat in the 21st century [1]. Specifically, there are several types of superbugs (bacterial strains resistant to all known antibiotics), including *A. baumannii, P. aeruginosa*, and *Enterobacteriaceae*, that have been listed as highest-priority targets for the future antibiotic drugs by the World Health Organization [2]. These bacteria are resistant to the most effective classes of conventional antibiotics available, including the broad-spectrum carbapanems: imipenem, meropenem, ertapenem, and doripenem [3,4]. It is known that these antibiotics target the inter-cellular environment and deactivate the enzymes that inhibit cell death, leading to the destruction of the bacteria cell (autolysis). They are meant to be administered as a last resort to preserve their efficacy, minimize toxicity and avoid the development of bacteria resistance. However, frequent exposure of bacteria to these antibiotics has already led to the development of resistance via mutations that stimulated the production of different enzymes to replace the deactivated autolysis inhibitor enzymes. Other resistance mutations are associated with creation of new efflux pumps to push the antibiotics out of the cell, and also include the porin mutations that prevent the antibiotics from penetrating the cell walls [3,4].

Antimicrobial peptides (AMPs), which include several classes of short and mostly positively-charged peptides, have been suggested as possible alternatives or adjuncts to conventional antibiotics [5]. Their positively-charged groups bind efficiently to negatively-charged segments of bacterial membranes, after which they stimulate the formation of pores in the cell walls. These pores eventually lead to the cell deaths [6]. AMPs play a critical role in the innate immune system as the first line of defense against invading pathogens, but they can also potentially serve as the most effective last line of defense [7]. Furthermore, two or more types of AMPs in combination, and also one type of AMPs in combination with an antibiotic, have shown the potential to be highly effective drugs with lower toxicity than the individual AMPs or antibiotics [8]. Remarkably, a single AMP type that effectively halts the bacterial growth can continue to exert significant antimicrobial activity when used in lower concentrations in combination with an ineffective AMP or resistant antibiotic, essentially sensitizing the bacteria to the antibiotic to which they were previously resistant [9]. Moreover, combinations of AMPs can remain effective in spite of bacteria evolution in cases in which bacteria would evolve resistance to individual AMPs in the combination [10].

While decades of experimental studies have tested multiple AMP-AMP and AMP-antibiotic combinations against the antibiotic-resistant bacteria, it is realistically not feasible to test quickly all possible combinations against all possible bacteria. The process from drug production and discovery to testing and releasing into the market is resource-costly, which is why various machine-learning methods, including recommendation systems, have been utilized to accelerate the identification of promising drug candidates [11–13]. With machine-learning methods, the existing data on AMP antimicrobial activity can be leveraged to narrow the pool of AMP combinations that need to be tested [14,15]. However, the main challenge is that when considering individual bacteria, the data on AMP and AMP-antibiotic efficacy are highly sparse. The data are even sparser for AMP-AMP combinations, which may contribute to the lack of studies on elucidating molecular mechanisms of antibacterial action for existing AMP-AMP combinations and predicting activity of new ones [16]. Previous studies focused on the efficacy of single AMP types alone and were limited in combining very different bacteria for their analyses (e.g., broad categories of gram-positive vs gram-negative or active against one type vs inactive against all tested types [17,18]). Although certain AMP types and AMP combinations have shown broad-spectrum antimicrobial activity, there are key differences between bacteria, for example in membrane composition [19] or morphology [20], that can affect the efficacy of a single AMP type or AMPs in combination and render some bacteria but not others resistant to the antimicrobial agents. It is important to develop new AMP-based antibiotic drugs by taking into account the specificity of the target bacterial species.

One method that takes into account such specificity is the user-based recommendation system, which was originally developed for predicting unknown movie ratings for one user based on a similar user who watched and rated those movies [21]. In particular, a recommendation-system method called neighborhood-based collaborative filtering (NBCF), has been successfully applied in biomedical research to predict protein-protein interactions [22], new anti-cancer drugs [23], drug side effects [24], and diseases for which existing drugs haven’t been tested and could potentially be effective [25]. NBCF seemed to be an ideal method also for investigating AMP antimicrobial activity across numerous bacteria because similarities in responses of different bacteria to the same AMPs can be used to predict responses of bacteria to an untested AMP type or AMP combination. One critical advantage of NBCF over other methods is that the minimum sparsity requirement for the data set is low, particularly as the number of users (bacteria) and items (AMP types or AMP combinations) increases. Moreover, the accuracy of the predictions is user-independent and data-dependent, so as the available amount of data on AMP efficacy increases, NBCF will be more useful and can provide reliable predictions that could not be made previously for a greater number of bacteria. The predictions of NBCF can then be fed to additional machine-learning models to augment the size of the training data and increase the prediction accuracy for aspects in which NCBF is limited [26].

In this work, we adopted the user-based NBCF method and applied it to predict antimicrobial activity of individual AMP types and AMP-AMP and AMP-antibiotic combinations. We found high predictive accuracy for known activity ratings for numerous bacteria. Our predictions can be tested since specific quantitative estimates that can be measured in experiments are obtained. Furthermore, we visualized AMPs and antibiotics as nodes in a network with edges representing synergistic or non-synergistic combined activity and predicted future edges representing AMP-Antibiotic and Antibiotic-Antibiotic combinations using a link prediction algorithm. We further investigated the synergistic combination activity of AMPs using their physico-chemical descriptors. The propy package [27] was utilized to extract the most important physico-chemical features of AMPs [18], and the similarity of two AMPs was defined as the inverse of their Euclidean distance in the space of these features. Strikingly, using our estimates of the similarities for different systems, we predict that the highest synergy in combinations of peptides is expected for those AMPs that are not too similar and not too dissimilar. Possible physico-chemical arguments to explain these optimal similarity observations are presented.

## Material and Methods

### Fractional Inhibitory concentration (FIC)

Antimicrobial activity can be evaluated through a parameter known as minimum inhibitory concentration (MIC), which is defined as the minimum concentration of an antimicrobial required for a population of bacteria to stop growing [28]. Synergy between antimicrobials, including AMPs and antibiotics, is commonly tested through a checkerboard assay in which the concentration of one antimicrobial is kept constant while the other is increased until the combination is effective in inhibiting bacteria growth, as presented in Fig. 1 [29].

**Fig 1.**
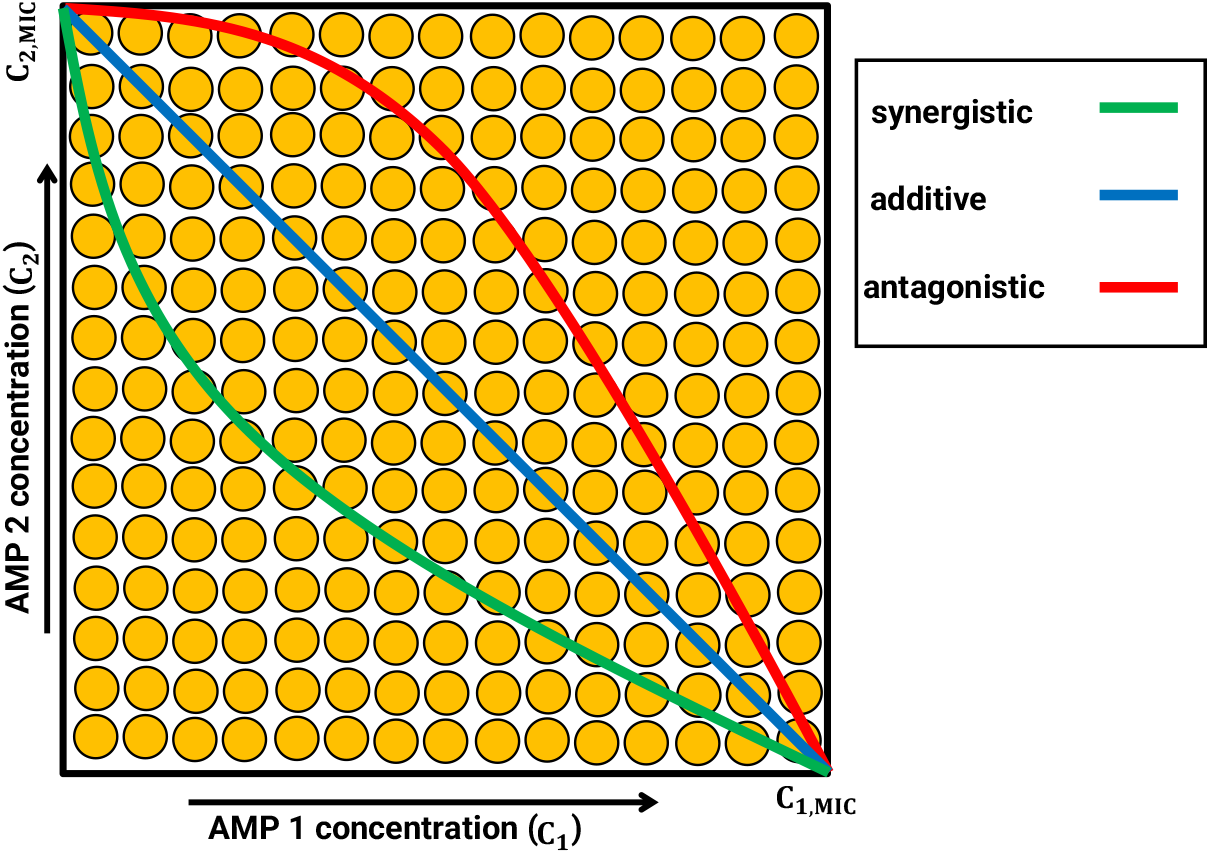
Schematic representation of synergy assay checkerboard for measuring cooperativity of antimicrobial combinations. The concentrations of AMPs of types 1 and 2 increase in *x* and *y* directions, respectively. The minimum inhibitory concentration (MIC) of AMP of type 1 (2) in the absence of AMP type 2 (1) is denoted as *C*_1,*MIC*_ (*C*_2,*MIC*_).

One of the ways to quantify the synergy between antimicrobial peptides is through evaluation of fractional inhibitory concentration (FIC) coefficients. For two antimicrobial agents, including an AMP and an antibiotic, labeled as 1 and 2, acting individually or in combination, the FIC indexes are defined as,

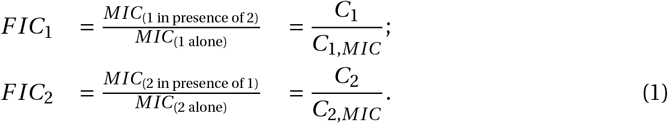

The combined FIC index is then given by *F IC = F IC*_1_ *+F IC*_2_ The antibiotic efficiency of the AMP combination is quantified then by the value of the combined FIC index as follows: *F IC <* 1 indicates the synergism, *F IC =* 1 corresponds to additivity, and *F IC >* 1 indicates the antagonism. Note, however, that for practical applications a stricter condition with *F IC ≤* 0.5 is typically applied to identify the strongest synergistic combinations of antibiotic drugs.

### Data Set Structure

We extracted data from the DBAASP database [30]. We filtered out AMPs that had synthetic amino acids or sequence length shorter than 11 to meet criteria for physico-chemical descriptor analysis in propy (propy analyzes natural amino acids and requires a length greater than 11 to calculate certain descriptors). Since predictions in bacteria-based collaborative filtering are made through nearest neighbors, the dataset must contain ratings for the same AMPs/combinations for at least two bacteria. Accordingly, we selected AMPs with reported MIC values for at least two bacteria, and if there was more than one MIC value for the same bacterium, we averaged all available MIC values to arrive at a single value per bacterium. Similarly, we selected AMP-Antibiotic and AMP-AMP combinations with corresponding FIC values for at least two bacteria, and if there were multiple FIC values for the same combination and bacterium, we calculated the mean and considered that as the FIC value for the bacterium in response to the AMP combination. To meet criteria for collaborative filtering, we removed columns (representing bacteria) and rows (representing AMPs) if they had fewer than two entries. We also removed bacteria columns that did not have any overlapping AMP/combination ratings with other bacteria columns. Thus, a bacterium was excluded from the data set if there were fewer than 2 AMP combinations targeting that bacterium, and similarly AMP combinations were excluded if they were tested on less than 2 bacteria. Bacteria were removed for not having ratings for at least two of the same AMPs/combinations as any other bacteria.

The sparsity of the ratings matrix is an important parameter in our analysis, and it is defined as:

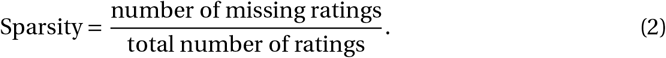

The ratings are MIC or FIC values depending on the system. To reduce data sparsity, we separated bacteria by species and collapsed across phenotypes within species, averaging ratings between phenotypes for the same AMP-combination. The final dataset included 12 bacteria and 49 pairs for AMP-AMP combinations; 49 bacteria and 50 pairs for AMP-antibiotic combinations, and 151 bacteria and 1040 AMPs for individual AMPs. A summary of the collected data, including for AMP-AMP and AMP-antibiotic combinations and individual AMPs, is shown in Fig. 2. Clearly, there are many missing values.

**Fig 2.**
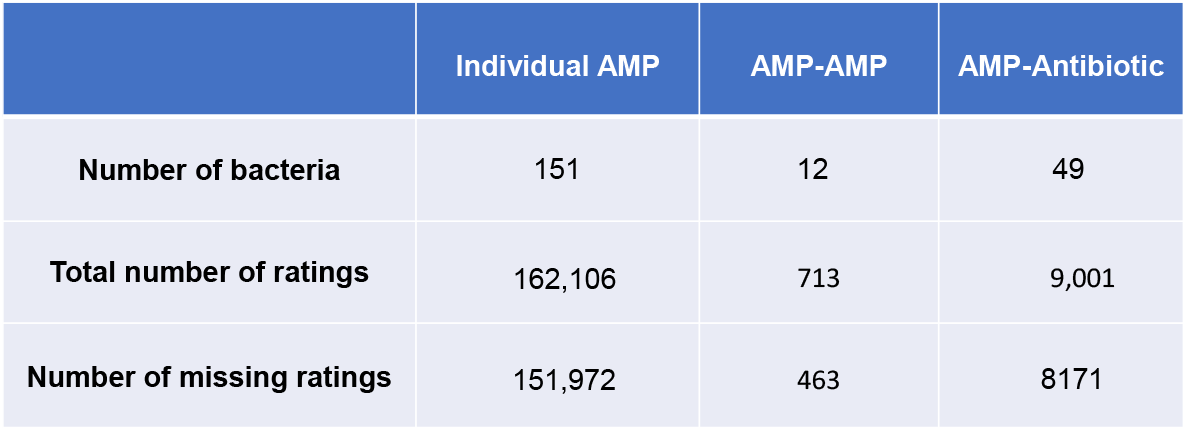
Summary of the data for individual AMPs, AMP-AMP combinations, and AMP-Antibiotic combinations. Number of bacteria refers to the number of distinct bacteria in each dataset. Number of missing ratings refers to the number of unknown antimicrobial activity values in each dataset. Total number of ratings reflects the dimensions of each dataset, equivalent to the number of distinct bacteria multiplied by the number of distinct AMPs or AMP combinations.

### Nearest-neighbor based collaborative filtering

For a given bacterium, depending on their patterns of ratings for AMP combinations, we can consider several nearest neighbors in the *d*-dimensional space (as visualized in Fig. 3a). The nearest neighbors for each bacterium can be determined using a proper similarity measure, which is a function that quantifies the similarity between two objects. Then, a threshold can be selected such that only pairs with similarity at or exceeding the threshold are considered nearest neighbors. For example, a correlation greater than 0.90 is implemented in our analysis, but in principle any value of the threshold can be used. We opted to use a Pearson correlation coefficient (*r*), which is an important measure that captures the similarity between the properties of two bacteria *α* and *β* by quantifying the correlations [21]. It is given by

**Fig 3.**
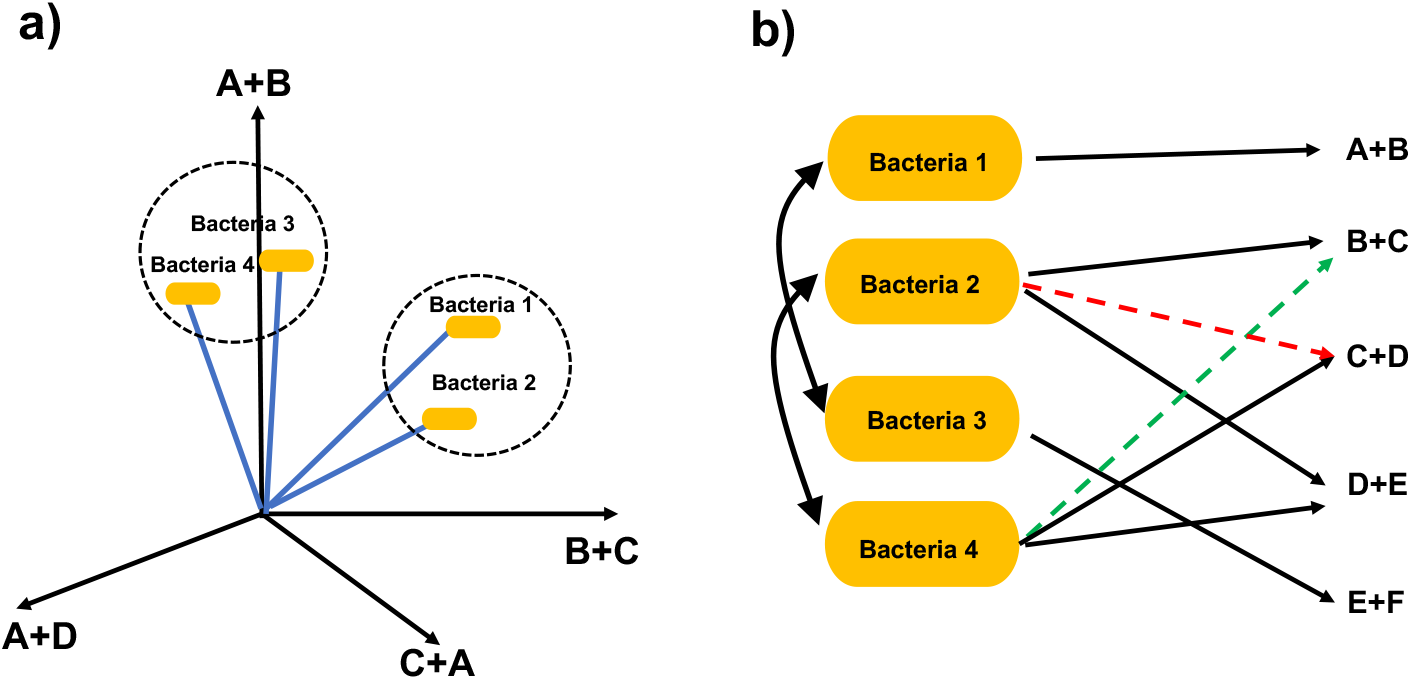
A schematic view of the nearest-neighbor based collaborative filtering approach. a) Each bacterium can be viewed as a point in d-dimensional space where each dimension represents an AMP or AMP combination (AMP-AMP or AMP-Antibiotic). b) Bacteria-based collaborative filtering. Bacterium 2 and Bacterium 4 are nearest neighbors because they are both inhibited by AMP combination D+E. Thus, based on similarity between Bacteria 2 and 4, we can predict the rating for Bacterium 2 in response to AMP combination C+D (antagonistic, red dashed line) and the rating for Bacterium 4 in response to AMP combination B+C (synergistic, green dashed line)

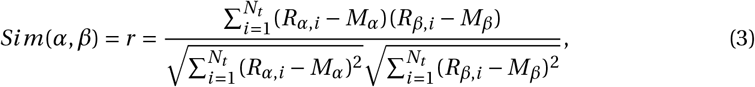

where *R*_*j, i*_ is the rating of combination *i* for bacteria *j*, and *M*_*j*_ is the average rating for bacteria 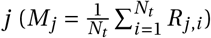, and *N*_*t*_ is the total number of available ratings. Using this similarity function, we can predict the unknown ratings for a given bacterium [21],

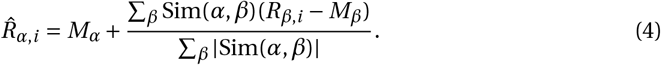

To test the quality of our predictions, we opted to use two quantitative measures. First measure is the coefficient of determination denoted as *R*^2^, which is defined as:

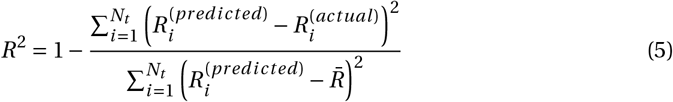

where 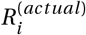 and 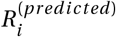 are actual and predicted ratings for AMP *i*, respectively, for a given bacterium. Also, 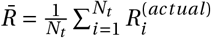 is the average of the actual ratings for a given bacterium. The closer *R*_2_ to unity, the better is prediction.

As a second measure of predictive accuracy, we use a root-mean square error (RMSQ), otherwise known as the quadratic mean, which is defined as follows,

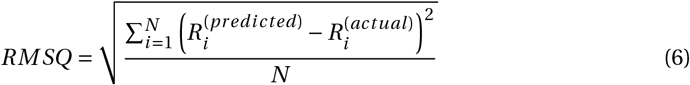

*R*^2^ is a standard method of evaluating statistical models, especially regression [31], and RMSQ is a suggested standard for evaluating specifically the recommender systems such as the collaborative filtering [32]. Higher *R*^2^ reflects greater predictive accuracy, and lower RMSQ reflects lower error. Alternatively, we can utilize normalized mean absolute error (NMAE), which can be used to compare datasets with different scales. This is particularly important because the values of FIC (describing the AMP-AMP combination) and MIC (describing individual AMP) have different ranges. NMAE is defined as

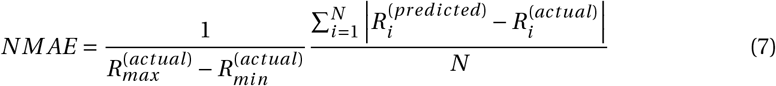

There are several advantages of applying the neighborhood-based collaborative filtering for our dataset. First, the properties of synthetic peptides cannot be analyzed by established AMP properties extractors, and therefore these properties cannot be used to predict antimicrobial activity. Because the user-based recommendation systems rely on similarities in activity between users (bacteria), synthetic peptides can be included in the analysis and their antimicrobial activity values can be predicted. Second, the antimicrobial activity values may differ according to species and strains of bacteria. While AMP antimicrobial activity has been determined for a wide range of bacteria, the AMP antimicrobial activity values for individual bacteria, when considered separately, are limited. Previous studies in their analysis frequently combined across the species of bacteria without taking into account the differences in membranes and other bacterial characteristics that could affect their antimicrobial activity. Our theoretical method allows us to estimate the antimicrobial activity for *specific* bacterial species.

### Link prediction

In this section, we propose to utilize a link method for predicting the possible synergistic AMP-AMP or AMP-Antibiotic combinations targeting specific bacteria [33]. This approach is particularly useful because *N* unique AMPs can be paired into *N* (*N −* 1)/2 unique combinations. To proceed further, let us visualize the rating pattern of a given bacteria to different AMP-Antibiotic combinations as a network, in which each AMP can be represented as a node and an edge between two nodes would indicate their synergistic combination. A snapshot of the corresponding network is visualized in Fig. 4. The full network for bacteria *E. coli* is shown in Fig. S1. In our data set, only a small subset of possible AMP-antibiotic combinations has been tested, and we would like to predict the potential synergistic combinations. We focused on *E. coli* since the link prediction must be conducted for one bacteria at a time. There were 80 unique AMPs and 15 unique antibiotics, and there were 92 AMP-Antibiotic pairs that are known to be synergistic.

**Fig 4.**
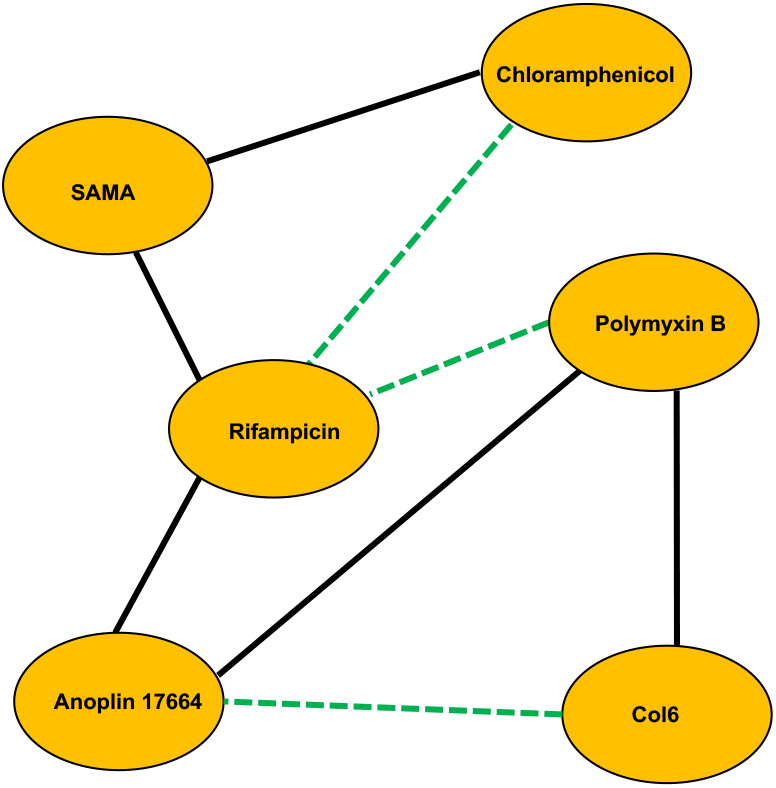
A schematic representation of the network of synergistic AMP-AMP combinations for a specific bacterium. Solid black lines connecting two nodes indicate that the corresponding AMPs synergize, while dashed green lines suggest potential synergistic AMP-AMP pairs. It is more likely that two AMPs will synergize if they have a common neighbor.

The link prediction method aims to determine the possibility of probable links in an incomplete graph. There are different methods for prediction of most probable links. Two widely used methods for link prediction are common neighbors and preferential attachments. Let us consider an arbitrary network in which two nodes *x* and *y* are not linked. We define *N* (*x*) and *N* (*y*) as a set of neighbors of node *x* and *y*, respectively. Then common neighbors of node *x* and node *y* is given by the intersection of the two sets:

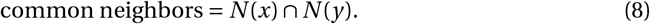

An alternative method for link prediction is the preferential attachment, which is defined as the multiplication of the number of neighbors of the node *x* by the number of neighbors of the node *y*,

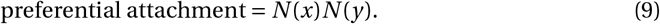

Therefore, each possible link between any two nodes can be characterized by the above characteristics. A link with highest score would be considered as most probable link in the complete network.

## Results

### Predicting individual AMP activity

In our analysis, the ratings for individual AMPs are minimum inhibitory concentration (MIC) values of antimicrobial activity against each bacterium. With 10,134 available ratings and 151,972 missing ratings, the sparsity of the data set was 0.94. The predictive accuracy, as shown in the scatterplot of 2060 actual vs predicted values from all bacteria in Fig. 5b, was high, *R*^2^*=* 0.92, NMAE = 0.01, but with a small number of large errors, RMSQ = 282.16.

**Fig 5.**
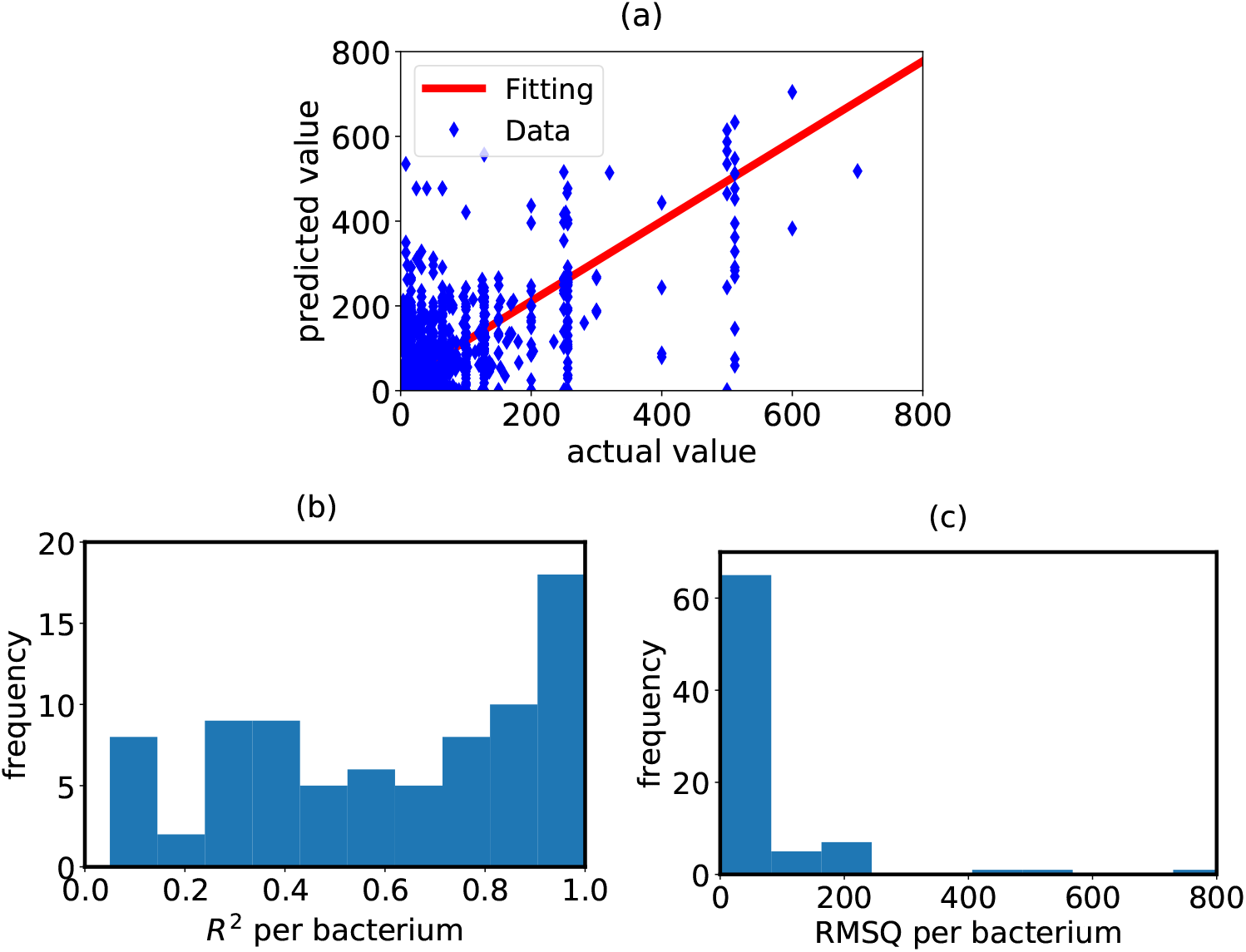
Collaborative filtering prediction validation for individual AMP antimicrobial activity. Scatterplot of 2060 actual vs predicted values, test subset for model validation, with line of best fit, *R*^2^ *=* 0.92, RMSQ = 28216, NMAE = 0.01. b) Histogram of *R*^2^ for individual bacteria. c) Histogram of RMSQ values for individual bacteria.

Specifically, the highest-frequency *R*^2^ range was 0.90-1, but the variability in *R*^2^ values across bacteria was high, as reflected in the long left-skewed tail of the distribution (see Fig. 5b). The distribution of RMSQ values in Fig. 5c suggests that most bacteria had values between 0 and 100, but there were bacteria with values ranging from 100 to 800, a small number of bacteria had large errors with high variability.

Table S1 shows additional relevant information, whihc is briefly summarized here. The number of bacteria nearest neighbors in the calculations of predicted scores ranged from 1 to 28, and most correlations between nearest neighbors were close to 1 (0.98 or higher). The number of data points in the calculation of *R*^2^ and RMSQ for each bacterium ranged from 2 to 372, for *S. aureus*. The number of predictions that could be made for each bacterium ranged from 2 to 943. The variability in parameters could be partially responsible for the variability in the predictive accuracy across bacteria as reflected in distributions of *R*^2^ and RMSQ.

### Predicting AMP-AMP combination activity

In our analysis, the ratings for AMP-AMP combinations are FIC values of antimicrobial activity against each bacterium. With 250 available ratings and 463 missing ratings, the sparsity of the data set was 0.65. The predictive accuracy, as shown in the scatterplot of 17 actual vs predicted values from all bacteria in Fig 6 (a), was satisfactory, *R*^2^ *=* .73, NMAE = 0.15, RMSQ = 0.31, but with a small number of bacteria (*n=* 5) for which predictions could be made. The distributions of *R*^2^ and RMSQ across bacteria in Figs 6 (b) and (c), respectively, show that though bacteria showed different *R*^2^ and RMSQ values, the variability was in a small range compared to corresponding distributions for individual AMPs.

**Fig 6.**
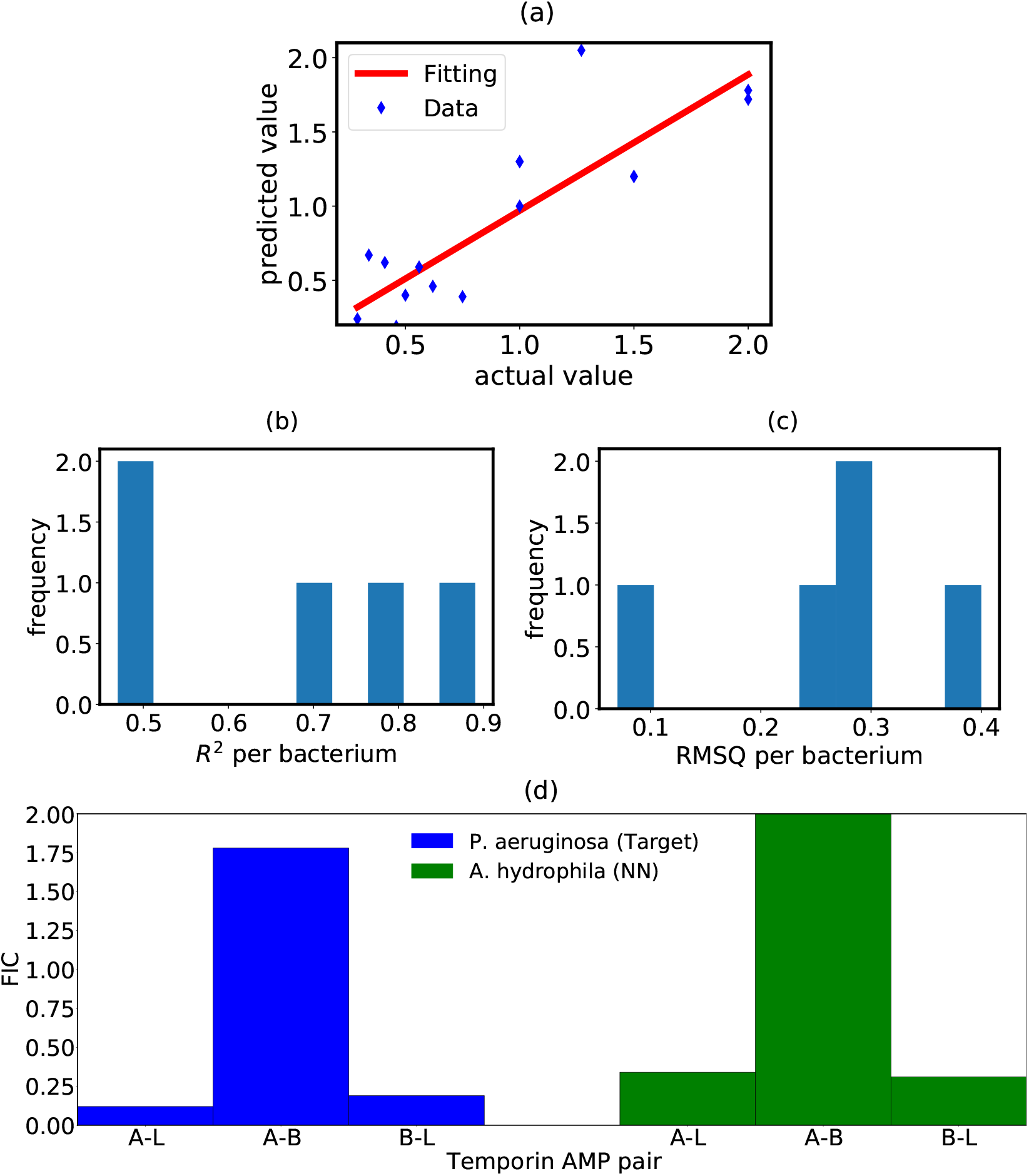
Collaborative filtering predictions validation for AMP-AMP antimicrobial activity. a) Scatterplot of 17 actual vs predicted values with the line of best fit, *R*^2^ = 0.73, RMSQ = 0.31, NMAE = 0.15. b) Histogram of *R*^2^ for individual bacteria. c) Histogram of RMSQ values for individual bacteria. d) Predicted FIC values for the bacterium *P. aeruginosa* (target) based on the nearest neighbor (NN) bacterium *A. hydrophila* for specific AMP-AMP pairs from the Temporin family (Temporins A, B, and L).

Table S2 shows additional relevant information: the minimum number of data points per bacterium for calculation of *R*^2^ and RMSQ was 3 and the maximum was 6, between 3 and 48 predictions could be made per bacterium, and the minimum correlation coefficient between nearest neighbors was 0.93. Because there were at most four nearest neighbors per bacterium for predictions, they were listed. The list of nearest neighbors shows that predictions could be made from one bacterium for another and vice versa. For example, *Staphylococcus aureus* was one of the nearest neighbors for *Bacillus subtilus* (*R*^2^ = 0.79), *Aeromonas hydrophila* (*R*^2^ = 0.47), and *Enterococcus faecalis* (*R*^2^ = 0.71), and these bacteria were also used to make predictions for *Staphylococcus aureus* (*R*^2^ = 0.49), though the prediction accuracy was not equally high for all bacteria. Notably, most of the antibiotic-resistant bacteria from the WHO priority list for which predictions could be made showed high predictive accuracy: *Acinetobacter baumannii* (*R* = 0.85, RMSQ = 12.99), *Enterobacter cloacae* (*R* = 0.80, RMSQ = 26.07), *Neisseria gonorrhoeae* (*R* = 1.0, RMSQ = 66.34), *Helicobacter pylori* (*R* = 1.0, RMSQ = 77.09), with the highest RMSQ for *Staphylococcus aureus* (*R* = 0.96, RMSQ = 240.22), and the lowest *R*^2^ for *Enterococcus faecium* (*R* = 0.43, RMSQ = 85.91).

As a specific example of how NBCF could be used to predict unknown values, the model evaluation for *P. aeruginosa* showed the highest predictive accuracy compared to that of other bacteria, *R*^2^ = 0.89, RMSQ = 0.25 (see Table S2). The nearest neighbor of *P. aeruginosa* was *A. hydrophila*, with the Pearson’s coefficient *r =* 0.94, and three FIC values could be predicted based on the similarity between the bacteria. As shown in panel d) of Fig. 6, the same AMP can be strongly or weakly synergistic depending on the second AMP in the combination.

Based on a threshold of *F IC ≤* 0.5 for the strong cooperativity between AMPs, the peptide Temporin A is predicted to be strongly synergistic with the Temporin L (FIC = 0.12) but strongly antagonistic with the Temporin B (FIC = 1.78). Temporin L, however, is predicted to synergize with both Temporin B (FIC = 0.19) and Temporin A. These predicted FIC values can be compared to the actual FIC values of the nearest neighbor, *A. hydrophila*, which show a similar pattern: Temporin L synergizes with Temporins A (FIC = 0.34) and B (FIC = 0.31), while Temporin A does not synergize with Temporin B (FIC = 2).

### Predicting AMP-Antibiotic combination activity

The ratings for AMP-antibiotic combinations were FIC values of antimicrobial activity against each bacterium. With 830 available ratings and 8171 missing ratings, the sparsity of the data set was 0.91. The predictive accuracy, as shown in the scatterplot of 116 actual vs predicted values from all bacteria in Fig 7 (a), was satisfactory, *R*^2^*=* .87, NMAE = 0.08, RMSQ = 0.20, and the distributions of *R*^2^ and RMSQ across bacteria revealed consistently satisfactory predictive accuracy: most of the bacteria had *R*^2^ values above 0.75 and RMSQ values below 0.1. Most of the predicted data points were close to the actual values and the line of best fit. As shown in Table S3, between 2 and 69 points points could be predicted, 2-23 points were used for *R*^2^ and RMSQ calculation, and there were 1-4 nearest neighbors per bacterium. Nearest neighbor correlation coefficients were above *r* = 0.92.

**Fig 7.**
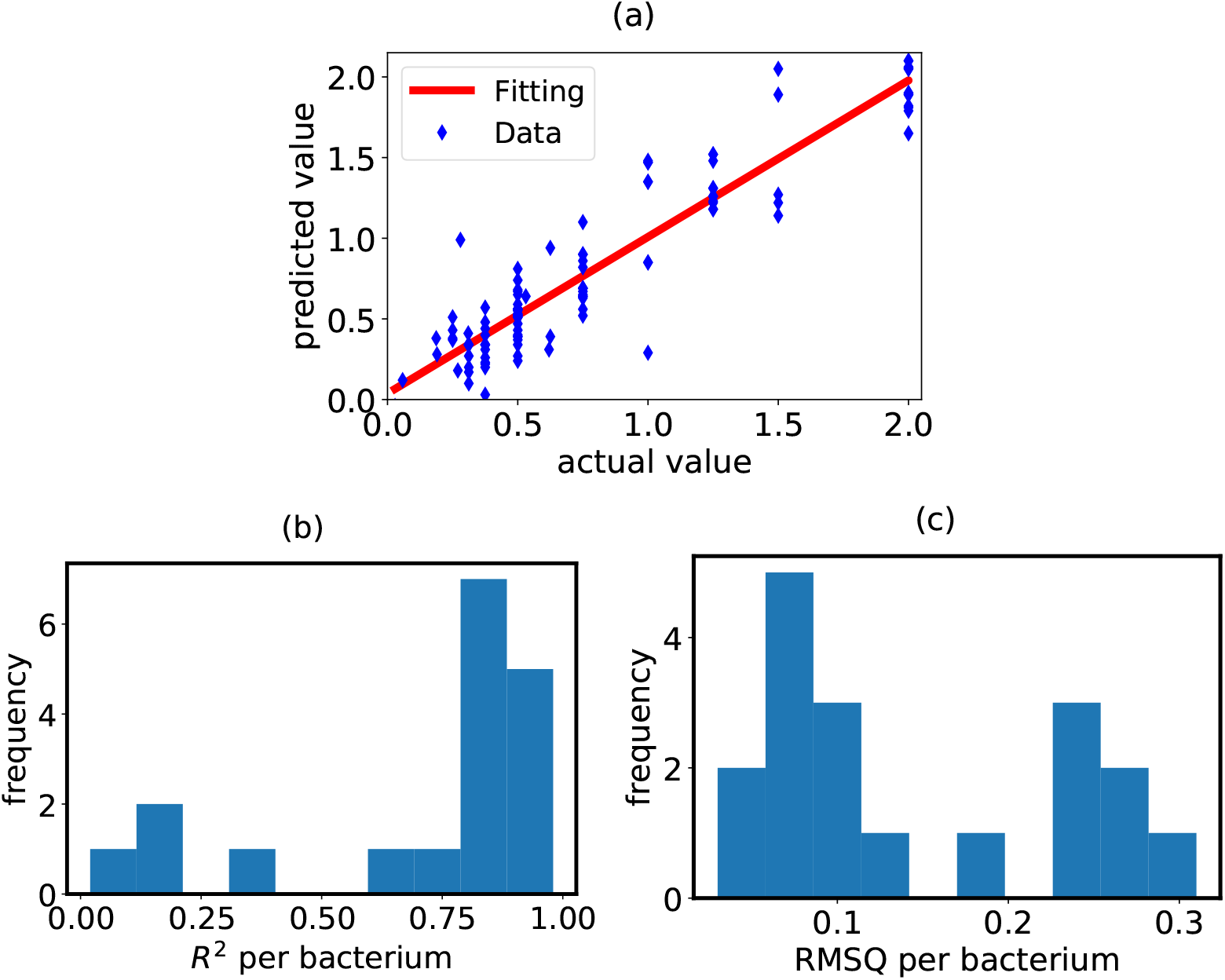
Collaborative filtering prediction validation for AMP-Antibiotic antimicrobial activity. a) Scatterplot of 116 actual vs predicted values with line of best fit, *R*^2^ = 0.87, RMSQ = 0.20, NMAE = 0.08. b) Histogram of *R*^2^ for individual bacteria. c) Histogram of RMSQ values for individual bacteria.

### Comparing NBCF results between datasets

A summary of the predictive accuracy across datasets is shown in Fig 8. The lower RMSQ for AMP-AMP and AMP-Antibiotic combinations could reflect that the subsets of data used for model evaluation are smaller and more homogeneous in prediction errors, while the individual AMP subset is larger and more heterogeneous. The NMAE takes into account the bias on the overall predictive accuracy coefficient due to predictions with large error that influences the RMSQ and shows that overall, errors were lowest for the individual AMP subset. The NMAE better reflects the *R*^2^ coefficient, and together, NMAE and *R*^2^ suggest that most predictions were accurate.

**Fig 8.**
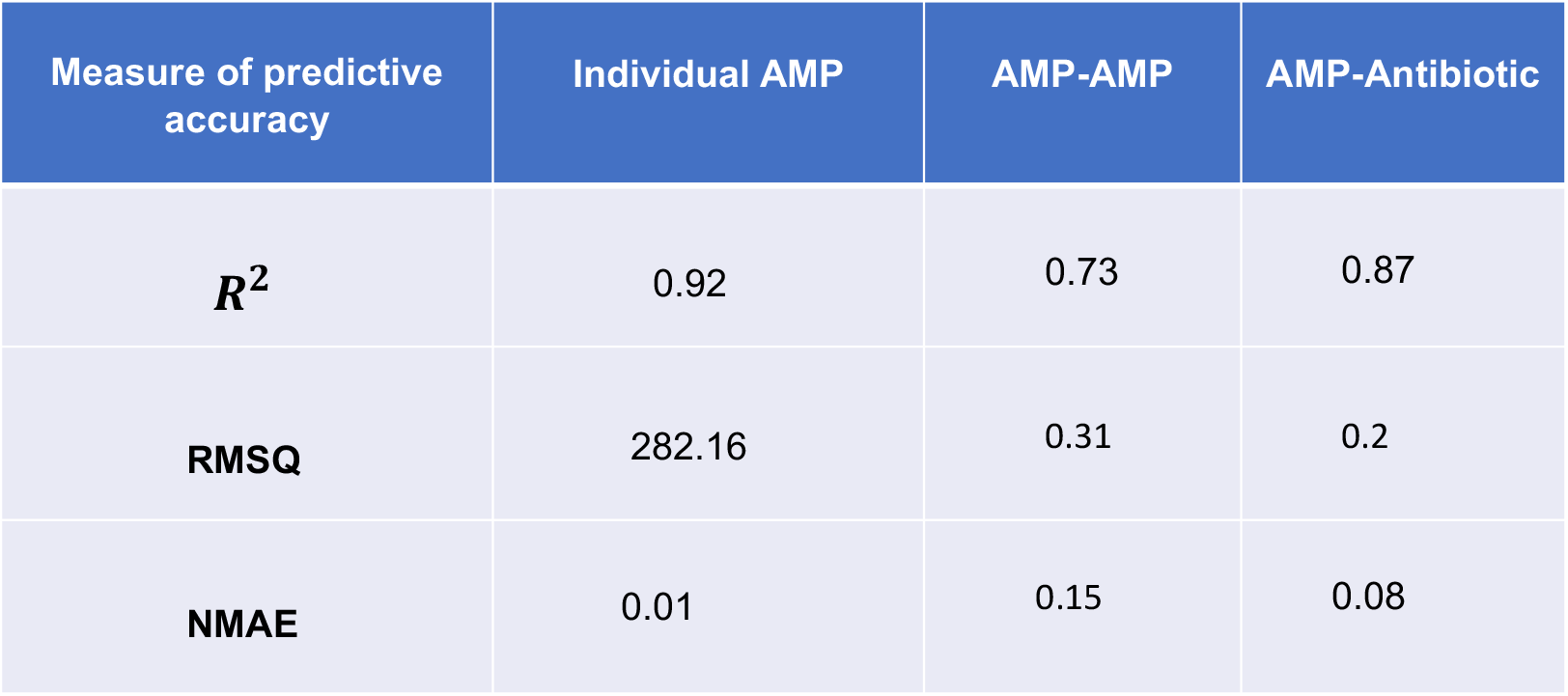
Comparison of different measures of predictive accuracy for collaborative filtering results.

### Predicting possible synergistic and non-synergistic AMP-Antibiotic pairs using the link method

The full network of synergistic AMP-antibiotic pairs is shown in Fig. S1. The largest number of AMP neighbors were for the antibiotics rifampicin (17), erythromycin (12), and kanamycin (8), and these antibiotics also had the largest number of common neighbors: 4 for rifampicin and kanamycin, 4 for rifampicin and erythromycin, and 5 for erythromycin and kanamycin. Thus, it is not surprising that by preferential attachment (see Fig. 9), antibiotics rifampicin and erythromycin, and rifampicin and kanamycin, had the highest likelihood of being synergistic and effective against *E. coli*. The method of preferential attachment also predicts the synergistic interactions between the rifampicin and the AMP Shiva 10 AMD and between the rifampicin and the AMP Col6. Polymyxin B, an AMP-based antibiotic, was predicted to synergize with the rifampicin and the erythromycin. We also predicted non-synergistic pairs from the network shown in Fig. S2. The AMP-based antibiotics vancomycin and polymyxin B were predicted not to synergize with each other or the non-AMP-based antibiotics gentamicin, rifampicin, and rifampin (see Fig. 9). Moreover, the AMPs with the sequence WRWRWR were predicted to be non-synergistic with the gentamicin.

**Fig 9.**
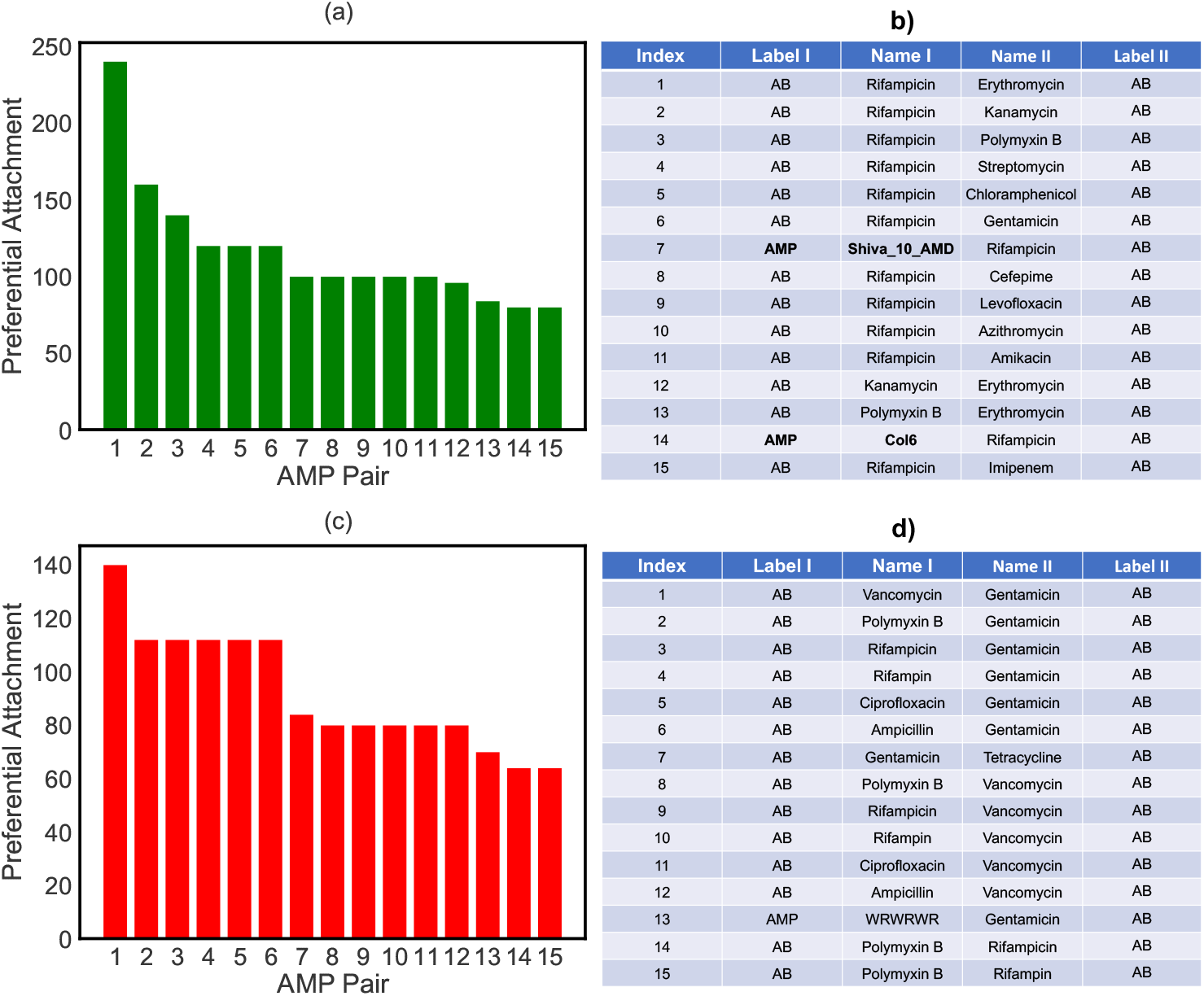
Likelihood of the predicted AMP-Antibiotic and Antibiotic-Antibiotic pairs targeting bacterium E. coli. by analyzing the preferential attachment for : a) Synergistic network b) A complementary table for the synergistic pairs. c) Non-synergistic network. d) A complementary table for the non-synergistic pairs.

### Elucidating the correlation between AMP combination activity and the similarity of two peptides

In recent years, different machine learning models have been developed to establish relationships between AMP sequence and biological activity. Since the predictability of successful machine learning model depends on the amount training data, building a structure-activity relationship for AMP combination might not work because of the very limited amount of available data. Yet, it is possible to elucidate the relationship between the physico-chemical features of two AMPs and their combination activity. The sequence of amino-acids by itself is not enough to fully describe the physico-chemical features of an AMP. From the peptide sequences, we can extract the physico-chemical features using available packages such as propy [27]. For each peptide, the package propy generates an ensemble of 1588 physico-chemical descriptors falling into five categories: basic character (e.g., charge), residue composition (e.g., dipeptide composition), auto-correlation, physico-chemical composition, and sequence order features.

Using the propy package descriptors, each AMP can be represented as a point in a high dimensional space. Thus the similarity of two peptides A and B is can be quantified by the Euclidean distance in this space:

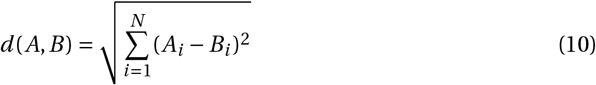

where *N =* 1588 is the number of descriptors. It can be argued that the similarity of two peptides is inversely proportional to this distance. In Fig. 10 we plotted the fractional inhibitory concentrations of 16 AMP-AMP combinations versus Euclidean distance. It can be seen that both similar pairs (distance *<* 9) and dissimilar pairs (distance *>* 11) are associated with greater values of FIC (*>* 0.5). However, for pairs with intermediate similarity (9 *<* distance *<* 11) the FIC values are less than 0.5. To make this observation more quantitative, we fitted a parabola (quadratic function) *F IC* (*A, B*) *= a*_2_*d* (*A, B*)^2^ *+a*_1_*d* (*A, B*) *+a*_0_ to describe these data. This result might indicate that there is an optimal value of similarly between AMP-AMP pairs that leads to strong synergistic interaction.

**Fig 10.**
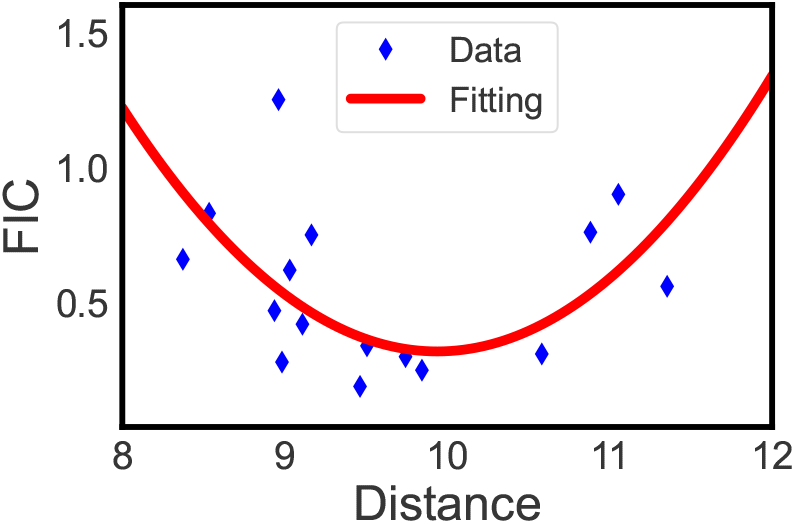
FIC coefficients of each AMP-AMP combination versus Euclidean distance between the peptides in pairs. A polynomial of degree 2 (quadratic function) *y = a*_2_ *x*^2^ *+a*_1_ *x +a*_0_ was fitted to the data. We utilized 16 AMP-AMP combinations targeting bacteria *E. coli*.

Although a peptide can be described by more than 1500 features, not all of these features are equally important. Indeed, only a limited number of important features determine the similarity of two given peptides. To test this hypothesis, let us consider the distance of two peptides in terms of an individual feature

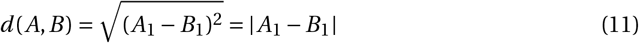

We analyzed the correlations between the similarities of two peptides in terms of each individual feature and corresponding FIC values for *E. coli* (Fig 11 (a)). Here we considered 12 most important features. A positive correlation between FIC and the distance means when two peptides are different in terms of a given descriptor, their combination activity is more likely to be antagonistic. The trend is opposite for negative correlations. An example is feature number 0 (MoreauBrotoAuto Hydrophobicity16), for which correlation is *−*0.63. In Fig 11 (c) we plotted FIC values versus distance (in MoreauBrotoAuto Hydrophobicity16) for a set of AMP-AMP combinations. It can be seen that dissimilarity of two peptides in terms of MoreauBrotoAuto Hydrophobicity16 leads to a synergistic combination.

**Fig 11.**
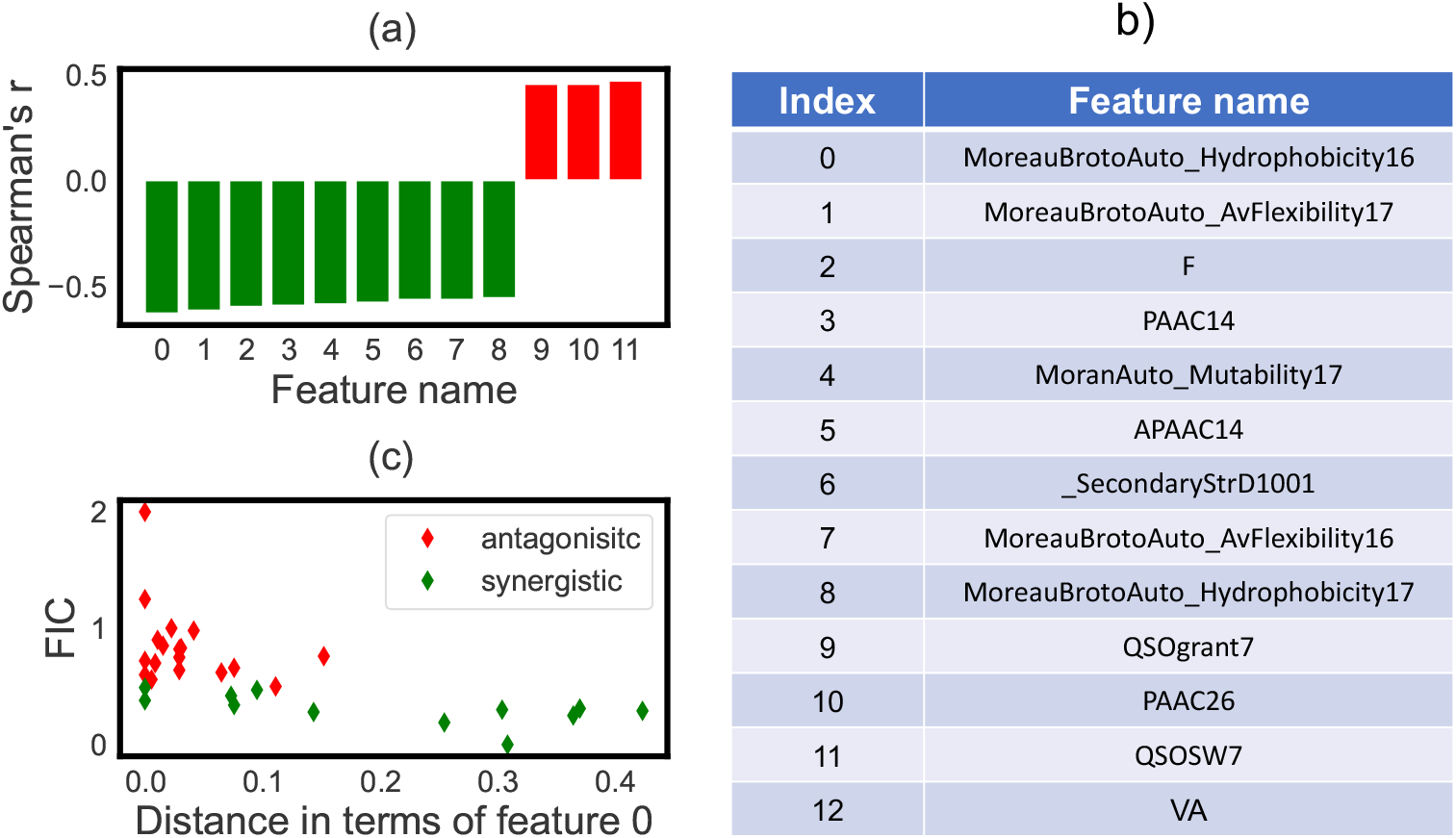
a) Spearman correlations (*r*) between FIC and distance of different AMP pairs in terms of specific descriptors. Green bars correspond to negative correlations, which suggests that dissimilarity of two peptides in terms of descriptors 1-4 leads to antagonistic combination. The trend is opposite for red bars. b) A complementary table for panel a. c) FIC versus distance in MoreauBrotoAutovHydrophobicity16 for synergistic (green) and antagonistic (red) pairs.

## Discussion

Using the method of nearest-neighbor collaborative filtering, we predicted the antimicrobial activity, quantified in MIC values, for bacteria in response to individual AMPs. The high predictive accuracy of the approach was shown in terms of *R*^2^, RMSQ, and NMAE for most bacteria, including the bacteria from the WHO priority list of antibiotic-resistant urgent threats, and we were able to predict values for these bacteria in response to untested AMPs. We also predicted the antimicrobial activity, quantified in FIC values, for the combinations of AMPs as well as for the combinations of AMPs and antibiotics that have not yet been reported for given bacterial species. Based on the identification of nearest neighbors for each bacterium using the Pearson’s correlation coefficients, the FIC parameters for several bacteria have been theoretically estimated. It is demonstrated for the antibiotic-resistant WHO priority target bacterium *P. aeruginosa* that the similarity with a less-threatening gram-negative bacterium, *A. hydrophila*, could be used to make predictions on untested AMP-AMP combinations. Like for *A. hydrophila*, The antimicrobial peptides Temporins L are predicted to synergize with Temporins A and B against *P. aeruginosa*, while Temporins A and B are predicted to be not cooperating at all. *P. aeruginosa* shows similarities with *A. hydrophila* in terms of the class as opportunistic pathogens [34], gram-stain (negative) [35], Type III secretion system (a target for developing antibiotics) [36], and they also have similar flagellum [37].

Nonetheless, NBCF may not necessarily select the nearest neighbors according to the structural similarity but rather by the functional similarity, specifically in response to AMPs. Two bacteria might be very different from structural or evolutionary point of view, but if their responses to a set of AMPs are similar because the AMPs can target both, they will be grouped by the NBCF as the nearest neighbors. This may present a danger in the accuracy of predictions because AMPs are very diverse and may target multiple species of bacteria or only one, and this might happen via different microscopic mechanisms. To probe the validity of our theoretical approach, it will be crucial to test our predictions with the quantitative experimental measurements for the same properties. The identification of effective AMP-antibiotic combinations is particularly important because antibiotics that have been expensive to develop can continue to be used in lower concentrations and with less toxicity and side effects when combined with the proper AMPs. Bacteria show reduced resistance to AMPs, AMP-AMP combinations, and AMP-antibiotic combinations in contrast to antibiotic-antibiotic combinations [10,38]. The benefit of AMP-antibiotic combinations is that even if a bacterium develops resistance to a certain combination, the same antibiotic can be used together with a different type of AMP, and the new combination may be effective with lower risk of the development of the resistance.

We also utilized the link prediction method to identify potential synergistic AMP-AMP pairs for a single bacterium, *E. coli*. We consider this method as a complementary approach to the nearest-neighbor collaborative filtering method since new combinations that are not yet established in experiments can be predicted. For example, pairs that have not yet been tested on any bacteria and thus have no FIC values from which to predict the unknown FIC values can be identified. While the bacterial-based collaborative filtering can extend results from one or more bacteria to predictions for another bacterium, link predictions can suggest new combinations that are likely to be effective but have not yet been tested on any bacteria. Moreover, if the collaborative filtering cannot be used to make predictions for a certain bacterium because there are no similar bacteria that can serve as nearest neighbors in the data set, the link prediction still can be used to make binary predictions (synergistic or not synergistic) and can potentially predict new AMP-AMP combinations and antibiotic-antibiotic combinations in addition to AMP-antibiotic combinations from a network that contains only AMP-Antibiotic edges. This method is especially important because it allows us to analyze the synergistic activities of AMP-AMP and AMP-antibiotic combinations for a specific bacterium.

An important factor in determining the synergistic activity of two different types of AMPs is a set of physico-chemical properties of the peptides. Using the package propy, we extracted these physico-chemical properties for different peptides that work in synergistic or non-synergistic pairs against *E. coli*. This allowed us to evaluate the similarity of two different peptides using the Euclidean distances in the corresponding multi-dimensional space. Our results on the similarity between AMPs in synergistic vs antagonistic pairs, using the propy-extracted descriptors, suggest that there is an optimal distance in features such as net charge and amino acid composition that corresponds to the highest synergy as measured by FIC. In other words, the strongest cooperativity between AMPs is expected when they are not too similar and not too dissimilar, so that there is some optimal similarity. This result is surprising and it might reflect the molecular details of the antimicrobial action of AMPs.

To understand these observations, it is important to briefly discuss the possible mechanisms of action by AMPs. It is currently widely believed that after the association to bacterial membranes AMPs stimulate the opening of the pores that break the cell walls and destroy the bacterial cells [39]. It is expected then that two different types of AMPs complement each other in the pore-opening process. For example, antimicrobial peptides PGLA and magainin-2 are highly synergistic, and their mechanism of action has been proposed to be through complementary functions: PGLA forms pores relatively fast, but these pores are not as stable as those formed by magainin-2 [40]. But together they can form fast and stable membrane pores. We argue that if two AMPs are too similar to each other they would not have enough different features to complement each other. But if AMPs are too dissimilar, they might not end at the same region of the bacterial cell membrane to exhibit any cooperativity. Thus, our theoretical analysis predicts that the highest synergy between AMPs is observed for some optimal similarity. It will be critically important to test this theoretical predictions in experiments.

AMPs is a diverse group of molecules that may target more than one bacterium effectively. They have been shown to be effective against a variety of pathogens and even against some diseases, including diabetes [41] and cancer [42]. Because some AMPs in the data sets that we utilized in our analysis show this multi-targeting ability and others do not, the bacteria-based collaborative filtering may have shown high predictive accuracy for some bacteria but not for others. Predictions will be more accurate for items in which nearest neighbors continue to have similar response patterns, however, for some AMPs the response patterns of two bacteria might diverge. For example, Table S1 for AMP-AMP combination collaborative filtering results showed that the predictive accuracy was not equally high across bacteria with similar nearest neighbors, potentially because for some AMPs predictions were less accurate. Collaborative filtering is data-dependent, and thus the predictions will be more reliable and accurate for larger and more homogeneous data sets. In the current study, the bacteria with greater data points for the calculations of *R*^2^ and RMSQ are likely to have more reliable estimates, and we recommend that these predictions to be tested first experimentally.

The observation that the prediction accuracy was not equally high for all bacteria also suggests that it is important to take into account the physico-chemical features of AMPs. Similarity of AMPs within the data set used for the collaborative filtering can be estimated through an analysis of AMP physico-chemical characteristics [43], as conducted in this study with the use of the propy package. Accordingly, future studies can use the analysis of physico-chemical characteristics to guide AMP-based collaborative filtering in addition to the bacteria-based collaborative filtering [44]. Furthermore, the current AMP prediction methods can be developed for the design of peptides for specific targets or the analysis of peptides discovered through RNA sequencing [45]. Physico-chemical descriptors of AMPs can be extracted with the propy package or similar approach, and machine learning models trained on AMP descriptors can generate predictions for specific bacteria [46], which can be expanded to all possible bacteria through the collaborative filtering. Finally, analyzing the physico-chemical descriptors that are most relevant for prediction through feature selection can help to develop microscopic models of AMP-AMP interactions and how to tune these interactions [47].

It is important also to discuss the limitations of our study. One of the biggest limitations is the size and the nature of the data sets. We expect that the link prediction and the collaborative filtering to become more relevant as more AMP-based drugs are tested on more bacteria, particularly on similar and overlapping bacteria. Specifically, it would be helpful for the same measure (e.g., FIC) to be reported, although other measures have been also proposed for AMP-antibiotic and AMP-AMP combinations tested in the same bacteria [48]. Another limitation is that we applied the models in their most simplified forms. Based on the demonstrated utility of the basic models for predicting AMP antimicrobial activity, more advanced models can be developed. For example, we can extract the latent features from the recommendation systems using matrix factorization [49], and we can investigate with more detail the bacterial similarity with the collaborative filtering based on the phylogenetic tree scores [50], a measure of evolutionary divergence, and the AMP similarity with the Boltzmann-machine-based multiple sequence alignment models [51]. All these directions will be investigated in the future.

## Author Contributions

A.M and H.T. designed the research. A. M. and H.T. performed the research. A. M and H.T. and A.B.K. wrote the article.

## Acknowledgments

The work was supported by the Welch Foundation (C-1559), by the NIH (R01 HL157714-02), by the NSF (CHE-1953453 and MCB-1941106), and by the Center for Theoretical Biological Physics sponsored by the NSF (PHY-2019745).

## References

1. O’Neill J. Tackling Drug-Resistant Infections Globally: Final Report and Recommendations-The Review on Antimicrobial Resistance. Date of access. 2016;16(09).

2. Tacconelli E, Carrara E, Savoldi A, Harbarth S, Mendelson M, Monnet DL, et al. Discovery, research, and development of new antibiotics: the WHO priority list of antibiotic-resistant bacteria and tuberculosis. The Lancet Infectious Diseases. 2018;18(3):318–327.

3. Suay-García B, Pérez-Gracia MT. Present and Future of Carbapenem-Resistant Enterobacteriaceae Infections. Advances in Clinical Immunology, Medical Microbiology, COVID-19, and Big Data. 2021; p. 435–456.

4. Aslam B, Rasool M, Muzammil S, Siddique AB, Nawaz Z, Shafique M, et al. Carbapenem resistance: Mechanisms and drivers of global menace. Pathog Bact. 2020;.

5. Magana M, Pushpanathan M, Santos AL, Leanse L, Fernandez M, Ioannidis A, et al. The value of antimicrobial peptides in the age of resistance. The Lancet Infectious Diseases. 2020;20(9):e216–e230.

6. Zhu Y, Hao W, Wang X, Ouyang J, Deng X, Yu H, et al. Antimicrobial peptides, conventional antibiotics, and their synergistic utility for the treatment of drug-resistant infections. Medicinal Research Reviews. 2022;.

7. Pasupuleti M, Schmidtchen A, Malmsten M. Antimicrobial peptides: key components of the innate immune system. Critical reviews in biotechnology. 2012;32(2):143–171.

8. Sierra JM, Viñas M. Future prospects for Antimicrobial peptide development: Peptidomimetics and antimicrobial combinations. Expert Opinion on Drug Discovery. 2021;16(6):601–604.

9. Hyun S, Choi Y, Jo D, Choo S, Park TW, Park SJ, et al. Proline hinged amphipathic α-helical peptide sensitizes gram-negative bacteria to various gram-positive antibiotics. Journal of Medicinal Chemistry. 2020;63(23):14937–14950.

10. Maron B, Rolff J, Friedman J, Hayouka Z. Antimicrobial Peptide Combination Can Hinder Resistance Evolution. Microbiology spectrum. 2022;10(4):e00973–22.

11. Li J, Tong XY, Zhu LD, Zhang HY. A machine learning method for drug combination prediction. Frontiers in Genetics. 2020;11:1000.

12. Güvenç Paltun B, Kaski S, Mamitsuka H. Machine learning approaches for drug combination therapies. Briefings in Bioinformatics. 2021;22(6):bbab293.

13. Jin W, Stokes JM, Eastman RT, Itkin Z, Zakharov AV, Collins JJ, et al. Deep learning identifies synergistic drug combinations for treating COVID-19. Proceedings of the National Academy of Sciences. 2021;118(39):e2105070118.

14. Wu L, Wen Y, Leng D, Zhang Q, Dai C, Wang Z, et al. Machine learning methods, databases and tools for drug combination prediction. Briefings in bioinformatics. 2022;23(1):bbab355.

15. Lee EY, Lee MW, Fulan BM, Ferguson AL, Wong GC. What can machine learning do for antimicrobial peptides, and what can antimicrobial peptides do for machine learning? Interface focus. 2017;7(6):20160153.

16. Nguyen TN, Teimouri H, Medvedeva A, Kolomeisky AB. Cooperativity in Bacterial Membrane Association Controls the Synergistic Activities of Antimicrobial Peptides. The Journal of Physical Chemistry B. 2022;.

17. Söylemez ÜG, Yousef M, Kesmen Z, Büyükkiraz ME, Bakir-Gungor B. Prediction of Linear Cationic Antimicrobial Peptides Active against Gram-Negative and Gram-Positive Bacteria Based on Machine Learning Models. Applied Sciences. 2022;12(7):3631.

18. Lee EY, Fulan BM, Wong GC, Ferguson AL. Mapping membrane activity in undiscovered peptide sequence space using machine learning. Proceedings of the National Academy of Sciences. 2016;113(48):13588–13593.

19. Oliveira J, Reygaert WC. Gram negative bacteria. 2019;.

20. Yang DC, Blair KM, Salama NR. Staying in shape: the impact of cell shape on bacterial survival in diverse environments. Microbiology and Molecular Biology Reviews. 2016;80(1):187–203.

21. Aggarwal CC. In: Neighborhood-Based Collaborative Filtering. Cham: Springer International Publishing; 2016. p. 29–70. Available from: https://doi.org/10.1007/978-3-319-29659-3_2.

22. Luo X, You Z, Zhou M, Li S, Leung H, Xia Y, et al. A highly efficient approach to protein interactome mapping based on collaborative filtering framework. Scientific reports. 2015;5(1):1–10.

23. Zhang L, Chen X, Guan NN, Liu H, Li JQ. A hybrid interpolation weighted collaborative filtering method for anti-cancer drug response prediction. Frontiers in pharmacology. 2018;9:1017.

24. Zhang W, Zou H, Luo L, Liu Q, Wu W, Xiao W. Predicting potential side effects of drugs by recommender methods and ensemble learning. Neurocomputing. 2016;173:979–987.

25. Newman R, Pietras CM, Qu F, Sapashnik D, Kofman L, Butze S, et al. Improving cell-specific drug connectivity mapping with collaborative filtering. bioRxiv. 2020;.

26. Aggarwal CC. Mining Web Data. In: Data Mining. Springer; 2015. p. 589–617.

27. Cao DS, Xu QS, Liang YZ. propy: a tool to generate various modes of Chou’s PseAAC. Bioinformatics. 2013;29(7):960–962.

28. Andrews JM. Determination of minimum inhibitory concentrations. Journal of antimicrobial Chemotherapy. 2001;48(Suppl_1):5–16.

29. Laishram S, Pragasam AK, Bakthavatchalam YD, Veeraraghavan B. An update on technical, interpretative and clinical relevance of antimicrobial synergy testing methodologies. Indian Journal of Medical Microbiology. 2017;35(4):445–468.

30. Pirtskhalava M, Amstrong AA, Grigolava M, Chubinidze M, Alimbarashvili E, Vishnepolsky B, et al. DBAASP v3: database of antimicrobial/cytotoxic activity and structure of peptides as a resource for development of new therapeutics. Nucleic acids research. 2021;49(D1):D288–D297.

31. Chicco D, Warrens MJ, Jurman G. The coefficient of determination R-squared is more informative than SMAPE, MAE, MAPE, MSE and RMSE in regression analysis evaluation. PeerJ Computer Science. 2021;7:e623.

32. Gunawardana A, Shani G, Yogev S. Evaluating recommender systems. In: Recommender systems handbook. Springer; 2022. p. 547–601.

33. Daud NN, Ab Hamid SH, Saadoon M, Sahran F, Anuar NB. Applications of link prediction in social networks: A review. Journal of Network and Computer Applications. 2020;166:102716.

34. Silvestry-Rodriguez N, Bright KR, Uhlmann DR, Slack DC, Gerba CP. Inactivation of Pseudomonas aeruginosa and Aeromonas hydrophila by silver in tap water. Journal of Environmental Science and Health, Part A. 2007;42(11):1579–1584.

35. Esselmann MT, Liu PV. Lecithinase production by gram-negative bacteria. Journal of Bacteriology. 1961;81(6):939–945.

36. Zhao YH, Shaw JG. Cross-talk between the Aeromonas hydrophila type III secretion system and lateral flagella system. Frontiers in microbiology. 2016;7:1434.

37. Wilhelms M, Molero R, Shaw JG, Tomás JM, Merino S. Transcriptional hierarchy of Aeromonas hydrophila polar-flagellum genes. Journal of bacteriology. 2011;193(19):5179–5190.

38. Sheard DE, O’Brien-Simpson Nm, Wade JD, Separovic F. Combating bacterial resistance by combination of antibiotics with antimicrobial peptides. Pure and Applied Chemistry. 2019;91(2):199–209.

39. Hollmann A, Martinez M, Maturana P, Semorile LC, Maffia PC. Antimicrobial peptides: interaction with model and biological membranes and synergism with chemical antibiotics. Frontiers in chemistry. 2018;6:204.

40. Chowdhury MH, Diamond G, Ryan LK. 11 Synergy of Antimicrobial Peptides. Antimicrobial Peptides: Discovery, Design and Novel Therapeutic Strategies. 2017; p. 188.

41. Soltaninejad H, Zare-Zardini H, Ordooei M, Ghelmani Y, Ghadiri-Anari A, Mojahedi S, et al. Antimicrobial peptides from amphibian innate immune system as potent antidiabetic agents: a literature review and bioinformatics analysis. Journal of Diabetes Research. 2021;2021.

42. Jin G, Weinberg A. Human antimicrobial peptides and cancer. In: Seminars in cell & developmental biology. vol. 88. Elsevier; 2019. p. 156–162.

43. Wang G. Bioinformatic analysis of 1000 amphibian antimicrobial peptides uncovers multiple length-dependent correlations for peptide design and prediction. Antibiotics. 2020;9(8):491.

44. Thakkar P, Varma K, Ukani V, Mankad S, Tanwar S. Combining user-based and item-based collaborative filtering using machine learning. In: Information and Communication Technology for Intelligent Systems. Springer; 2019. p. 173–180.

45. Moretta A, Salvia R, Scieuzo C, Di Somma A, Vogel H, Pucci P, et al. A bioinformatic study of antimicrobial peptides identified in the Black Soldier Fly (BSF) Hermetia illucens (Diptera: Stratiomyidae). Scientific reports. 2020;10(1):1–14.

46. Vishnepolsky B, Gabrielian A, Rosenthal A, Hurt DE, Tartakovsky M, Managadze G, et al. Predictive model of linear antimicrobial peptides active against gram-negative bacteria. Journal of chemical information and modeling. 2018;58(5):1141–1151.

47. Wang CH, Bashor CJ, Mehta P. The strength of protein-protein interactions controls the information capacity and dynamical response of signaling networks. arXiv preprint arXiv:181105371. 2018;.

48. Yu G, Baeder DY, Regoes RR, Rolff J. Combination effects of antimicrobial peptides. Antimicrobial agents and chemotherapy. 2016;60(3):1717–1724.

49. Zhang W, Yue X, Lin W, Wu W, Liu R, Huang F, et al. Predicting drug-disease associations by using similarity constrained matrix factorization. BMC bioinformatics. 2018;19(1):1–12.

50. Lalucat J, Mulet M, Gomila M, García-Valdés E. Genomics in bacterial taxonomy: impact on the genus Pseudomonas. Genes. 2020;11(2):139.

51. Gabler F, Nam SZ, Till S, Mirdita M, Steinegger M, Söding J, et al. Protein sequence analysis using the MPI bioinformatics toolkit. Current Protocols in Bioinformatics. 2020;72(1):e108.

